# Identification of locally activated spindle-associated proteins in oocytes uncovers a phosphatase-driven mechanism

**DOI:** 10.1101/2025.05.19.654801

**Authors:** Xiang Wan, Hiroyuki Ohkura

## Abstract

The meiotic spindle forms only around the chromosomes in oocytes, despite the exceptionally large volume of the cytoplasm. This spatial restriction is likely to be governed by local activation of key microtubule regulators around the chromosomes in oocytes, but the identities of these microtubule regulators and the mechanisms remain unclear. To address this, we developed a novel assay to visualise spatial regulation of spindle-associated proteins in *Drosophila* oocytes by inducing ectopic microtubule clusters. This assay identified several proteins including the TPX2 homologue Mei-38 that localise more strongly to microtubules near the chromosomes than away from them. We identified a microtubule-binding domain containing a region highly conserved also in humans. The domain itself is regulated spatially, and contains a conserved serine and a nearby PP2A-B56 docking motif. A non-phosphorylatable mutation of this serine allows the domain to localise to ectopic microtubules as well as spindle microtubules, while mutations in a PP2A-B56 docking motif greatly reduced the spindle localisation. As this phosphatase is concentrated at the kinetochores, it may act as a novel chromosomal signal spatially regulating spindle proteins within oocytes.

## Introduction

Due to the absence of centrosomes, the spindle is self-assembled in the oocytes of most animal species. As the oocytes have a large volume, it is crucial to assemble one bipolar spindle around the chromosomes and suppress spindle formation in other parts of the ooplasm. This spatially restricted spindle assembly requires the local activation of key proteins important for bipolar spindle assembly near the chromosomes. This spatial regulation is not only limited to microtubule nucleation factors but also to other regulators of microtubule dynamics and organisation, because non-spindle microtubules are present all over the oocyte yet fail to organise into a spindle. Chromosomes serve as spatial cues, which can be sensed by key regulators of the spindle assembly. These proteins can then be locally activated to execute their functions, together with many other ubiquitous proteins whose activity is not spatially regulated.

The Ran-Importin pathway is the most studied system for spatial regulation in oocytes. Ran is a small GTPase with two forms: active RanGTP and inactive RanGDP. RanGDP is converted into RanGTP by chromatin-bound RCC1, the guanine nucleotide exchange factor (GEF) [1]. Cytoplasmic Ran GTPase-activating protein (RanGAP) promotes GTP hydrolysis to GDP [2]. Due to chromosomal localisation of RCC1, the RanGTP concentration is higher around the chromosomes, generating a RanGTP gradient [3]. Proteins important for spindle assembly are inhibited by Importin binding, and RanGTP releases them from Importin for local activation near the chromosomes [4, 5]. These proteins are collectively called “spindle assembly factors”. So far, over 20 spindle assembly factors have been identified [6].

Among the spindle assembly factors, TPX2 is one of the first and most studied factors. TPX2 plays multifaceted roles in spindle assembly in meiosis. After RanGTP releases TPX2 from importin inhibition, TPX2 interacts with Aurora A to activate it and with Kinsein-5 to modulate its distribution at the spindle. TPX2 also forms condensates on microtubule lattices and recruits other microtubule nucleators, Augmin and γ-TuRC, for branching microtubule nucleation [7–10]. TPX2 plays crucial roles in spindle microtubule assembly, pole focusing and size control in mouse oocytes and/or *Xenopus* egg extract [11, 12].

In *Xenopus* egg extracts, a dominant negative Ran prevents spindle microtubule assembly, while hyperactive RanQ69L can induce spindle-like structures in the absence of chromosomes [13–16]. However, in mouse oocytes, abolishing the Ran-GTP gradient either by expressing dominant negative or hyperactive Ran leads to defects in the meiosis I spindle without disrupting spatially-restricted spindle assembly [17]. Similarly, in *Drosophila* oocytes, these dominant mutants affects spindle organisation, but a spindle is formed only around the chromosomes [18]. In human oocytes, expressing dominant negative Ran severely delayed impaired spindle assembly around the chromosomes [19]. Therefore, these observations suggest a varied importance of the Ran pathway in different species, but also the presence of alternative mechanisms for the spatial regulation of proteins important for bipolar spindle assembly in oocytes.

The chromosomal passenger complex (CPC) containing Aurora B kinase was proposed as an alternative signal for spatial regulation in oocytes. The CPC is essential for spindle microtubule assembly in *Xenopus* egg extract and in *Drosophila* oocytes [20–22]. The CPC is activated on the chromosomes independently of Ran [23], and directly phosphorylates and inactivates microtubule-destabilising proteins, such as MCAK/KIF2C [24–26] and Op18/Stathmin [27]. In addition, the CPC co-operates with the phospho-docking protein 14-3-3 to locally activate spindle-associated proteins around the chromosomes in *Drosophila* oocytes. 14-3-3 binds to Kinesin-14 Ncd to prevent it from binding to non-spindle microtubules away from the chromosomes. Chromosome-bound CPC phosphorylates an additional site on Kinesin-14 Ncd to release it from inhibition by 14-3-3 [28]. Many more spindle-associated proteins have been found to be regulated by 14-3-3, including the CPC and the centralspindlin complex [29].

Rcc1 and the CPC are two chromosomal cues so far identified for the spatial regulation of spindle-associated proteins in oocytes. However, it is unknown whether they can account for all of the spatial regulation in oocytes, as no systematic studies have been done to identify spatially-regulated spindle proteins, chromosomal signals or regulatory mechanisms. In this report, we developed a novel method to assess whether the binding of a protein to microtubules is spatially regulated by inducing ectopic microtubules in *Drosophila* oocytes. Our findings on the *Drosophila* TPX2 homologue Mei-38 suggest a protein phosphatase-based mechanism that spatially regulates this spindle-associated protein.

## Results

### Induction of ectopic microtubules revealed spatial regulation of Mei-38 binding to microtubules in *Drosophila* oocytes

To identify the spindle-associated proteins that are locally activated near the chromosomes and bind to microtubules, we have induced ectopic microtubule clusters by treating *Drosophila melanogaster* oocytes with a microtubule stabilising drug taxol/paclitaxel (**Fig 1A**). After taxol treatment, we immunostained wild-type mature oocytes using antibodies against various spindle-associated proteins or mature oocytes expressing GFP-tagged spindle-associated proteins using a GFP antibody, together with an α-tubulin antibody and DAPI.

**Figure 1.**
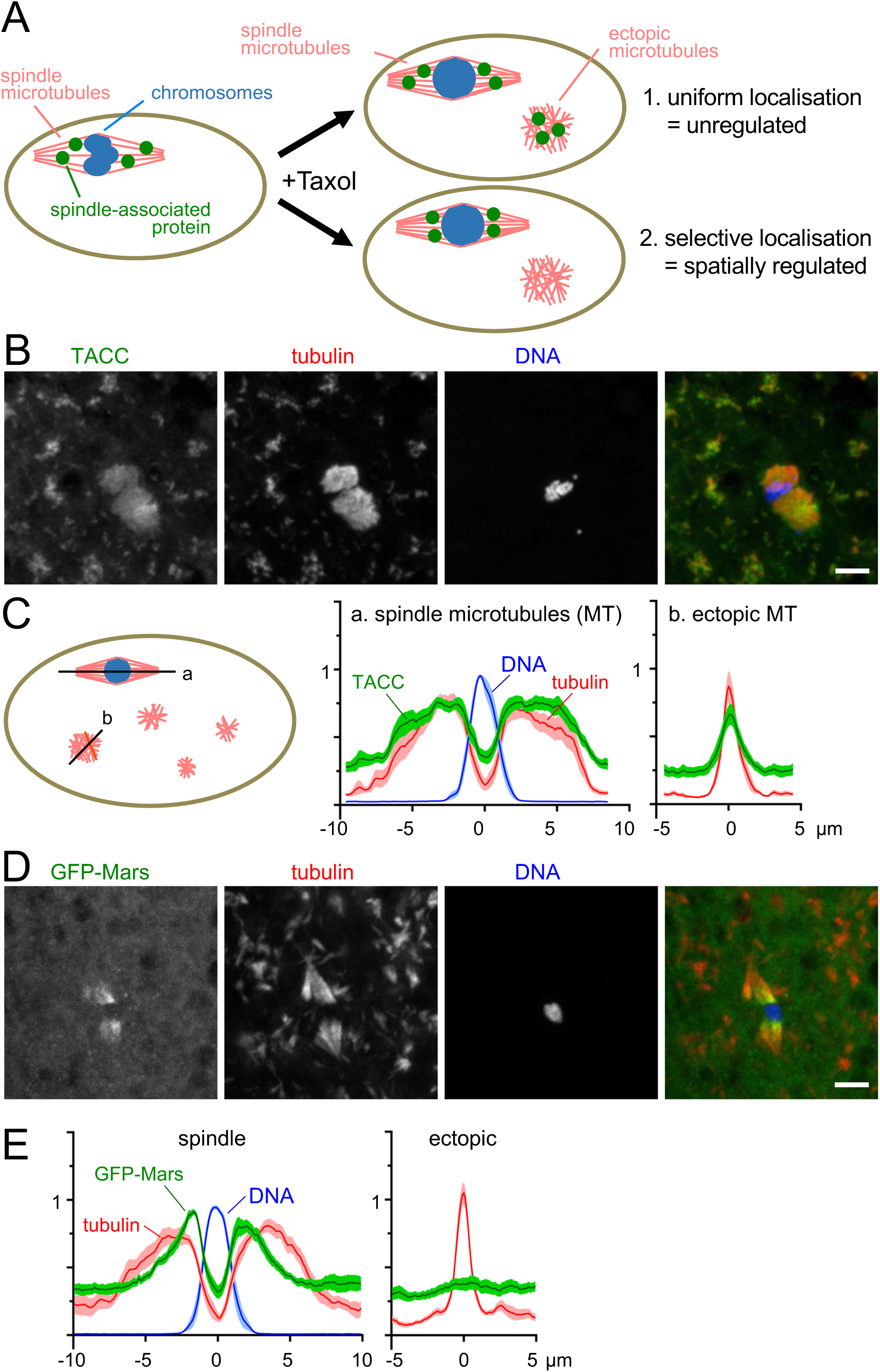
Induction of ectopic microtubules identified spindle-associated proteins that are spatially regulated in *Drosophila* oocytes. (A) Induction of ectopic microtubules by taxol reveals two types of spindle-associated proteins. Proteins that localise uniformly to spindle and ectopic microtubules are not spatially regulated in terms of microtubule binding. In contrast, proteins that selectively localise to spindle microtubules are spatially regulated. (B) TACC localisation in taxol-treated oocytes. Wild-type mature oocytes, which are naturally arrested in metaphase I, were incubated with taxol and immunostained using TACC and α-tubulin antibodies. TACC uniformly localised to both spindle and ectopic microtubules. Arrowheads indicate chromosomes. Bar=5 μm. (C) Quantification of TACC, α-tubulin and DNA signal intensities. The first line was drawn along the long axis of the spindle microtubules. The second line was drawn on an ectopic microtubule cluster with a similar maximum intensity to the spindle microtubules. The centre of the chromosomes or the ectopic cluster is defined as 0. Signal intensities were measured along the lines and normalised to the maximum intensity value of the spindle line in each oocyte. The graphs show means±s.e.m. (D) GFP-Mars localisation in taxol-treated oocytes. Mature oocytes expressing GFP-Mars were incubated with taxol and immunostained using GFP and α-tubulin antibodies. GFP-Mars localised to the spindle microtubules, not to ectopic microtubules. Within the spindle microtubules, the signal is stronger near the chromosomes. Bar=5 μm. (E) Quantification of GFP-Mars, α-tubulin and DNA signal intensities following the same method as C.

Without taxol treatment, mature *Drosophila* oocytes naturally arrest in metaphase I with a bipolar spindle. After a brief taxol treatment, the bipolar spindle associated with the chromosomes was still recognisable, although the morphology was often distorted to various degrees. In addition, many ectopic microtubule clusters were formed all over the oocytes (**Fig 1A**). The intensity and morphology varied but the maximum signal intensity of some clusters was comparable to that of the spindle microtubules. For the localisation of spindle proteins, we classified the outcomes into two types depending on whether they are spatially regulated (**Fig 1A**). (1) If a spindle protein indiscriminately binds to microtubules, it would equally localise to the entire spindle and ectopic microtubule clusters. (2) If a protein has a higher affinity to microtubules near the chromosomes than further away, a higher concentration of the protein would be observed on the spindle microtubules near chromosomes than ectopic microtubules with a similar microtubule density.

Among 12 spindle proteins that we tested, three showed equal localisation to all microtubules (**Fig 1A and S1 Fig**), indicating they are unregulated (type 1). They include TACC, KInesin-13 Klp10A, and Kinesin-5 Klp61F. For example, TACC showed uniform distribution on the spindle microtubules and ectopic microtubules, reflecting the intensity and pattern of tubulin signals (**Fig 1B**). To quantify the signal intensity, a line was drawn along the long axis of the spindle and another line was drawn over a microtubule cluster with roughly the same tubulin intensity as the spindle microtubules (**Fig 1C**). The intensity of the signal was measured along the lines, and normalised using the maximum intensity of the spindle microtubules in each oocyte. The mean signal intensity of TACC roughly follows the tubulin mean intensity along the spindle axis and also along the line drawn across ectopic microtubule clusters (**Fig 1C**). These proteins are likely to bind to microtubules indiscriminately under this condition. In other words, they are not regulated spatially in terms of microtubule binding.

In contrast, the other nine proteins showed stronger signals on the spindle microtubules, especially near the chromosomes, than ectopic microtubules indicating that they are spatially regulated (type 2). They include Mars/HURP, the CPC subunits, Ncd/Kinesin-14, Mink/NuSAP, Cyclin B and Mei-38/TPX-2, (**Fig 1C and S2 and 3 Fig**). For example, the signal of the GFP-tagged Mars was much stronger on spindle microtubules than on ectopic microtubule clusters, even where the tubulin intensity was comparable (**Fig 1D**). Furthermore, the GFP-Mars signal was stronger near the chromosomes, and became weaker away from the chromosomes even within the same spindle (**Fig 1D**). Quantification showed that the GFP-Mars signal intensity on the spindle microtubules decays much more sharply further away from the chromosomes than the tubulin intensity. Moreover, the GFP-Mars signal on ectopic microtubules is comparable to the background without a distinct peak (**Fig 1E**). Therefore, Mars binding to microtubules is strongest near the chromosomes, becomes progressively weaker further away along the spindle, and virtually absent on ectopic microtubules, indicating that Mars is spatially regulated in terms of microtubule binding.

### Truncation analysis narrowed the region sufficient for spatial regulation

We focused our study on Mei-38, the TPX2 homologue, among the spatially regulated proteins we identified, as it apparently lacks sequences potentially regulated by previously known mechanisms, such as importin-binding sites and 14-3-3 binding sites. Mammalian TPX2 is regulated by Ran-importin through nuclear localisation signals [4] but Mei-38 lacks nuclear localisation signals, as well as the Aurora A and Kinesin-5 interaction domains found in mammalian TPX2 ([30]; **Fig 2A**).

**Figure 2.**
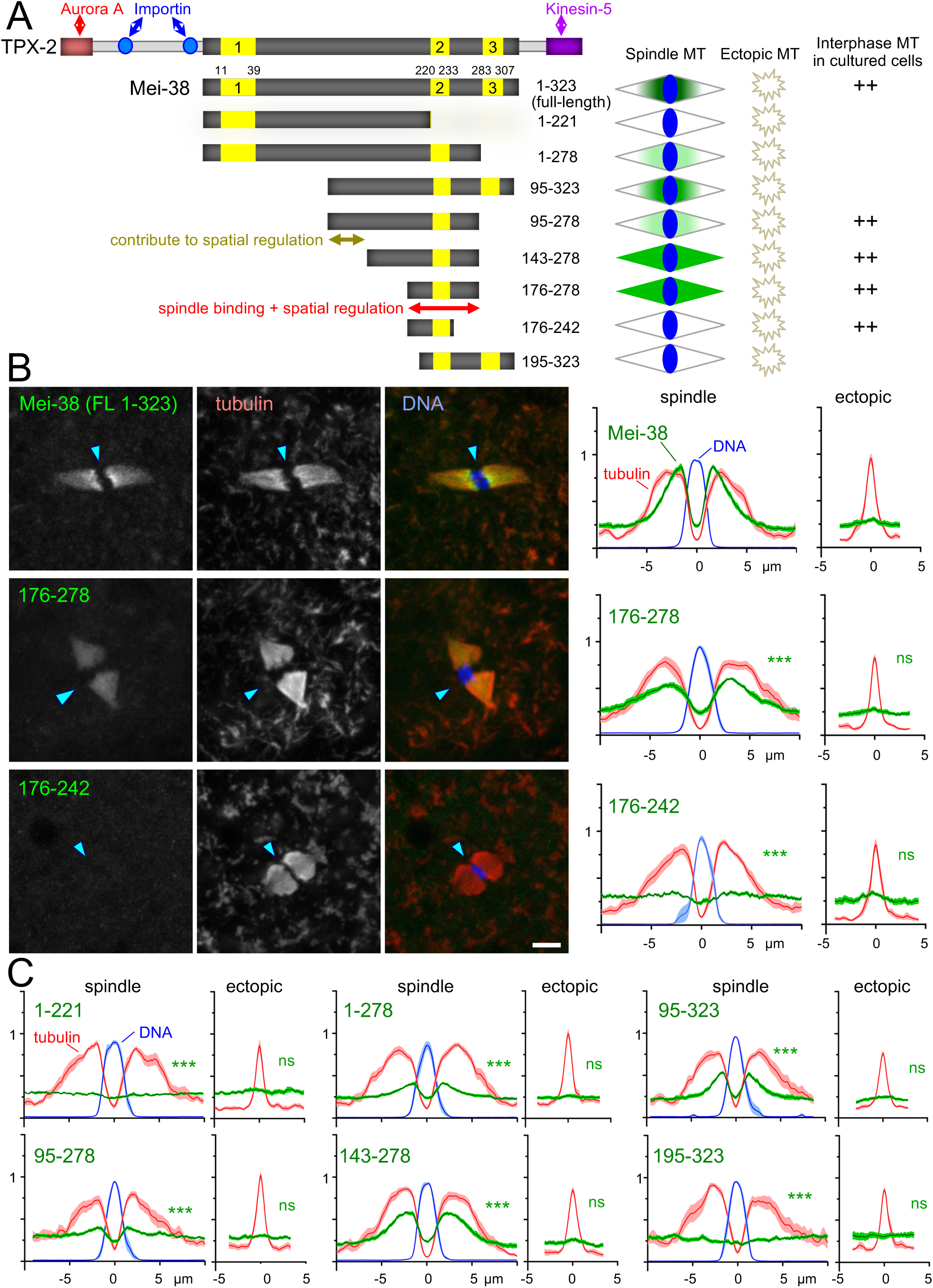
Truncations of Mei-38 defined a spindle-binding domain that is regulated spatially. (A) A summary diagram of Mei-38 truncations and their localisation in taxol-treated oocytes and interphase S2 cells. Numbers indicate amino-acid residues. The region including the conserved domain 2 is important for microtubule binding and spatial regulation. (B) Localisation of GFP-tagged Mei-38 truncations in taxol-treated oocytes. The arrowheads indicate the position of the chromosomes. Quantification were carried out as Fig.1C. For comparison, the profile of the Mei-38 mean signal intensity of each truncation is presented after normalisation to adjust the mean background signal intensity (at 0 in the spindle line) to be identical to the control (the full-length Mei-38). Bar=5 μm. *** and ns indicate p<0.001 and p>0.05, respectively, in two-tailed t-tests when the signal intensity of a GFP-Mei-38 variant is compared to that of GFP-Mei-38. (C) Quantification of the remaining truncations as Fig.2B. Images were shown in S5 Fig.

In taxol-treated oocytes, Mei-38 localised to the spindle microtubules much more strongly than to ectopic microtubule clusters, even where the tubulin intensity was comparable (**Fig 2B**. Furthermore, within the same spindle, the Mei-38 signal was stronger near the chromosomes and became gradually weaker towards the poles (**Fig 2B**). Quantification showed that the Mei-38 signal intensity on the spindle microtubules decays more sharply further away from the chromosomes than the tubulin intensity. Moreover, the Mei-38 signal on ectopic microtubules is close to the background with a very small peak (**Fig 2B**).

To define domains responsible for spindle binding and its spatial regulation, we carried out a taxol assay in oocytes expressing various GFP-tagged truncated Mei-38 proteins (**Figs 2A and 2B and S5 Fig**). Goshima (2011)[30] has identified three regions conserved among the TPX2 family across species including humans. Mei-38(1-278), Mei-38(95-323) and Mei-38(95-278), lacking the conserved region 1, 3 or both, still maintained spatial regulation identical to the full-length protein, although the signal intensity is lower than the full-length Mei-38 (**Figs 2A and 2C and S5 Fig**). This demonstrates that these regions are not essential for spindle localisation or spatial regulation.

Mei-38(143-278) and (176-278) localised uniformly to spindle microtubules but much more weakly to ectopic microtubules (**Figs 2A and C and S5 Fig**), suggesting that the area between the conserved regions 1 and 2 contributes to spatial regulation to some degree. Mei-38(1-221) lacking the conserved regions 2 and 3 failed to localise to any microtubules (**Figs 2A and 2C and S5 Fig**), suggesting the conserved region 2 and the C-terminal flanking area are important for microtubule association. We have identified Mei-38(176-278) containing the conserved region 2 as the shortest fragment sufficient to localise to spindle microtubules. This fragment is still under spatial regulation, as it showed much stronger localisation to the spindle microtubules than to ectopic ones. (**Fig 2B**). A smaller fragment, Mei-38(176-242), failed to localise to any microtubules. A western blot confirmed that this fragment was expressed at a similar level to the full-length or other truncated proteins (**S6 Fig**). Mei-38(195-323) failed to localise to any microtubules (**Fig 2C**), probably because of a very low protein level seen in a western blot (**S6 Fig**).

In summary, we narrowed down the region sufficient for the spindle localisation and spatial regulation in oocytes, although another region also contributes to spatial regulation.

### Identification of a microtubule-binding domain containing a highly conserved sequence

To assess the microtubule-binding activity separately from oocyte-specific regulation, we took advantage of a *Drosophila* embryonic cultured cell line, S2. A previous report (Goshima, 2011)[30] showed that GFP-Mei-38 expressed in S2 cells is associated with filamentous microtubule networks in interphase, and assessed the microtubule binding activity of some truncated proteins. We found that fragments that bind to spindle microtubules in oocytes, including Mei-38(176-278), were able to associate with interphase microtubules in S2 cells (**Fig 3A**). Unexpectedly, Mei-38(176-242), which failed to localise to any microtubules in oocytes, was able to associate with interphase microtubules in S2 cells (**Fig 3A**). Therefore, this deleted sequence 243-278 is dispensable for microtubule binding *in vitro* or in S2 cells, but required for spindle microtubule localisation in oocytes.

**Figure 3.**
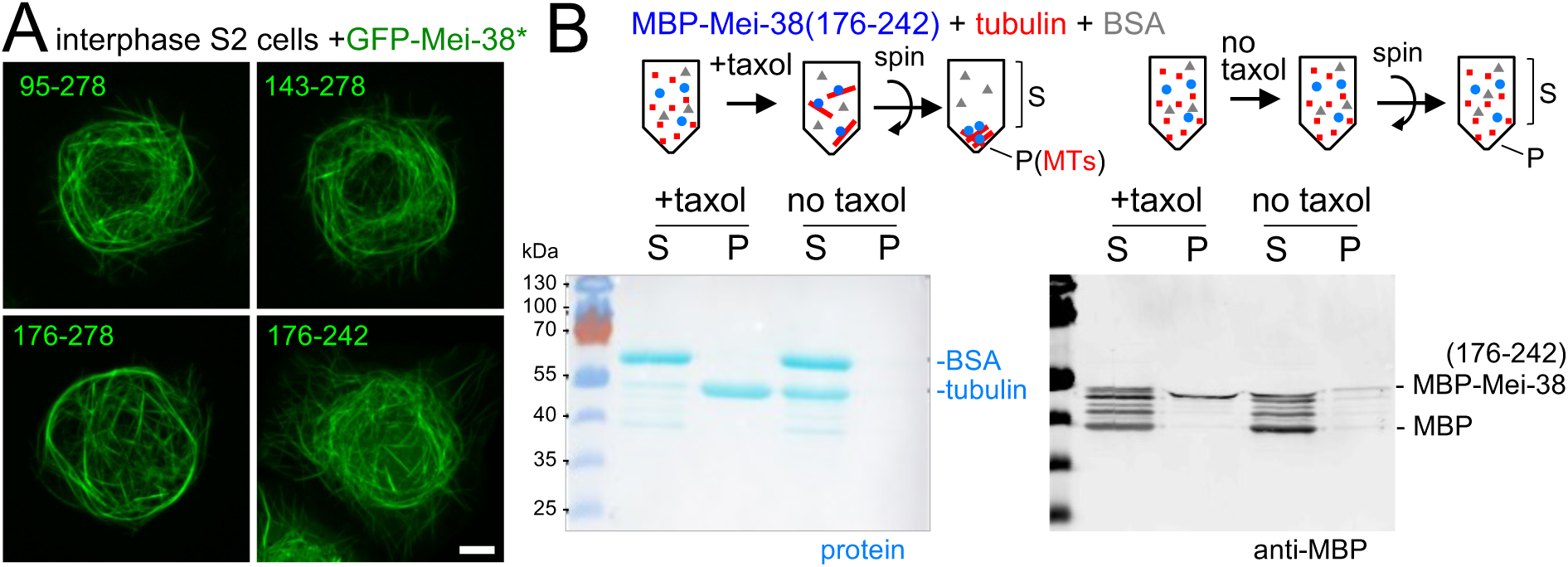
Mei-38(176-242) binds to microtubules in interphase S2 cells and *in vitro*. (A) Interphase S2 cells expressing GFP-tagged truncated Mei-38 known to associate with interphase microtubules. Mei-38(176-242) that does not localise to any microtubules in oocytes binds to microtubules in interphase S2 cells. Bars=5 μm. (B) Microtubule-binding assay of Mei-38(176-242aa). Purified MBP-Mei-38(176-242aa) expressed in bacteria was mixed with tubulin and BSA. Taxol was added to one sample (+taxol), but not to the other sample (no taxol). Microtubules (MTs) were polymerised in the sample with taxol, both samples were centrifuged. The supernatants (S) and pellets (P) were analysed by protein staining and western blot using an MBP antibody. Twice the amount of the pellets was loaded compared to the supernatants in each lane.

Mei-38(176-242) contains region 2, which is highly conserved also in vertebrates [30]. To test whether Mei-38(176-242) has intrinsic microtubule binding activity *in vitro,* we produced the fragment in bacteria as a fusion with MBP (**S7 Fig**). Pure porcine tubulin dimers were incubated with bacterially produced Mei-38(176-242) and were polymerised into microtubules by taxol addition. After microtubules were spun down, the microtubule fraction (pellet) and non-binding fraction (supernatant) were analysed by western blot using an MBP antibody (**Fig 3B**). MBP-Mei-38(176-242) was found in the microtubule fraction, while degradation products including the one with the size corresponding to MBP were missing (**Fig 3B**). In agreement with S2 cells, this showed that the recombinant Mei-38(176-242) has an intrinsic microtubule-binding activity *in vitro*. However, it fails to localise to the spindle or ectopic microtubules in oocytes.

### S225 is responsible for spatial regulation of the Mei-38(176-278) binding to microtubules

We then decided to focus on Mei-38(176-278), the smallest fragment that localises to the spindle in oocytes, because it is still spatially regulated. This fragment shows significantly stronger localisation to the spindle microtubules compared to ectopic microtubules in taxol-treated oocytes (**Fig 4C and D**). It contains three sequences of interest (**Fig 4A**): (1) a central highly conserved region containing the potential phosphorylation site S225 (**Fig 4B**), (2) the N-terminal flanking region 198-203 which matches a docking motif of PP2A-B56 (L/F/xxI/VxE) [31], (3) the C-terminal flanking sequence 243-278 that is dispensable for microtubule-binding *in vitro* or in S2 cells, but required for spindle localisation in oocytes.

**Figure 4.**
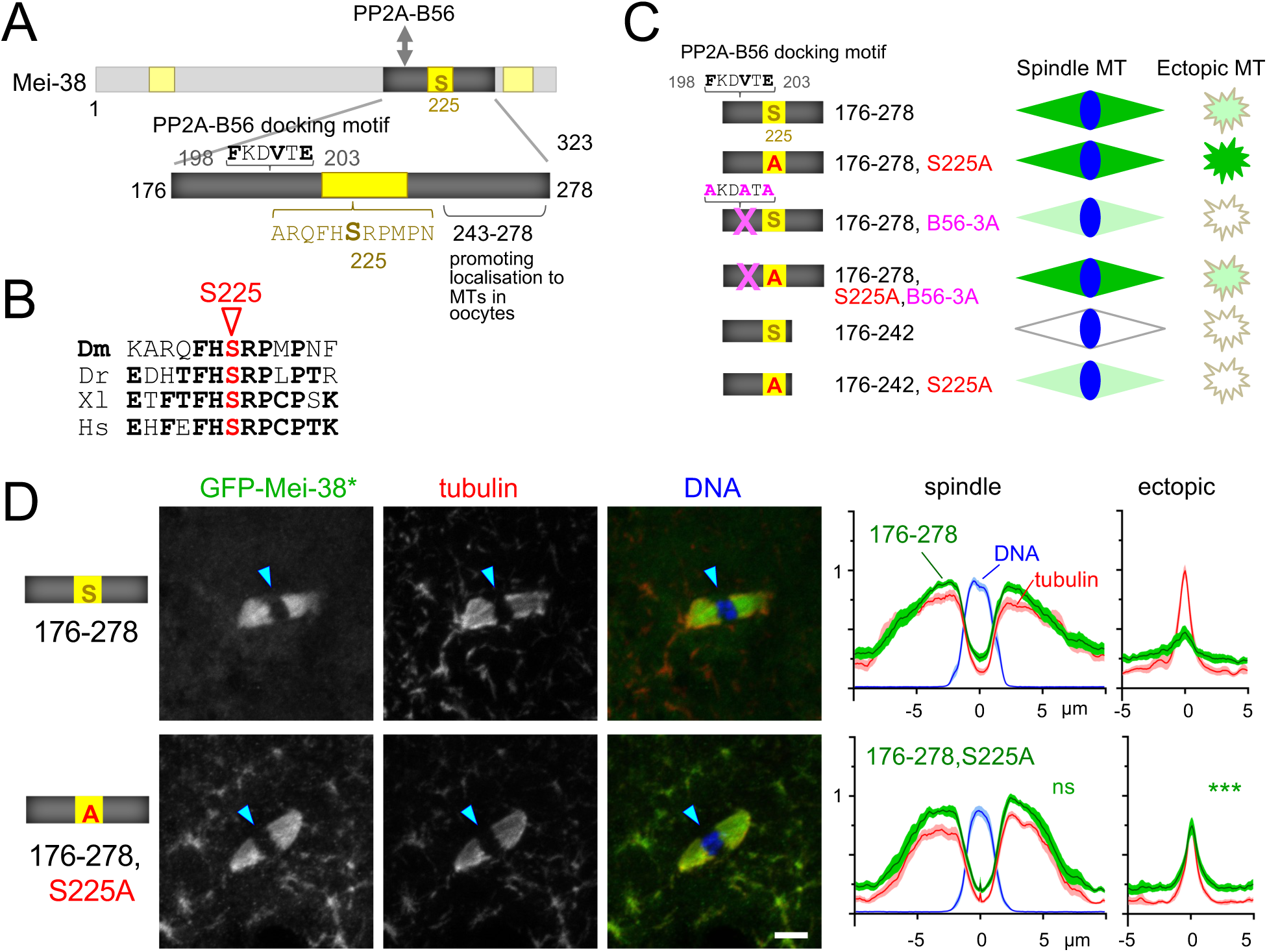
A non-phosphorylatable mutation (S225A) abolished spatial regulation of Mei-38(176-278) (A) The minimal fragment (176-278) under spatial regulation contains three regions of interest: a PP2A-B56 docking motif, the conserved region 2 with potential phosphorylation site (S225), and a region (243-278) dispensable for microtubule binding but important for spindle association in oocytes. (B) The conserved region 2 containing S225 is conserved also in vertebrates. Dm; *Drosophila melanogaster*. Dr; *Danio rerio*. Xl; *Xenopus laevis*. Hs; *Homo sapiens*. (C) A summary diagram of various mutants and truncations within the minimum fragment under the control of spatial regulation (176-278) with the localisation in taxol-treated oocytes shown in Fig.4 and 5. B56-3A; three key residues of the PP2A-B56 docking motif were mutated to alanine. S225A; a non-phosphorylatable mutation of S225 within the conserved region 2. 176-242; a region that can bind to microtubules in S2 cells and *in vitro*, but cannot localise to spindle microtubule in oocytes. (D) GFP-Mei-38(176-278) and GFP-Mei-38(176-278,S225A) localisation in taxol-treated oocytes. A non-phosphorylatable mutation at S225 (S225A) allowed a GFP-Mei-38(176-278) to localise to all microtubules equally, abolishing the spatial regulation. The arrowheads indicate the position of the chromosomes. Bar=5 μm. The signal distribution was quantified as Fig.1B. *** and ns indicate p<0.001 and p>0.05, respectively, in two-tailed t-tests when the signal intensity of a GFP-Mei-38 variant is compared to that of 176-278. For comparison, the profile of the Mei-38 mean signal intensity of each truncation is presented after normalisation to adjust the mean background signal intensity (at 0 in the spindle line) to be identical to Mei-38(176-278). Note that a microscope different from the one for Fig.1 and 2 was used to acquire the data in this figure and the following figures, and gave a lower background.

We tested the role of S225 in binding of Mei-38(176-278) to spindle and ectopic microtubules in taxol-treated oocytes by mutating it to alanine (S225A) (**Figs 4B and C**). Strikingly, this mutated fragment, Mei-38(176-278,S225A), localised to all microtubules equally including ectopic microtubules, completely abolishing spatial regulation of microtubule binding (**Fig.4D**). This result demonstrated that S225 is solely responsible for spatial regulation of this fragment.

### PP2A-B56 docking enhances Mei-38 localisation to spindle microtubules

Next we looked into the role of a flanking PP2A-B56 docking motif located near S225. This sequence is conserved among *Drosophila* species. To test its role, we mutated 3 consensus residues of this motif to alanines (B56-3A) in Mei-38(176-278) (**Fig 4A**). This mutation greatly reduced the localisation of Mei-38(176-278) to the spindle microtubules (**Fig 5A**). It suggests that a failure to dephosphorylate this fragment by PP2A-B56 suppresses the localisation to spindle microtubules. To test the involvement of S225 in this process, we further introduced a non-phosphorylatable mutation, S225A, to generate Mei-38(176-278,B56-3A,S225A). The spindle microtubule localisation was restored by S225A in taxol-treated oocytes (**Fig 5A**), suggesting that dephosphorylation of S225 by PP2A-B56 is important for spindle microtubule localisation.

**Figure 5.**
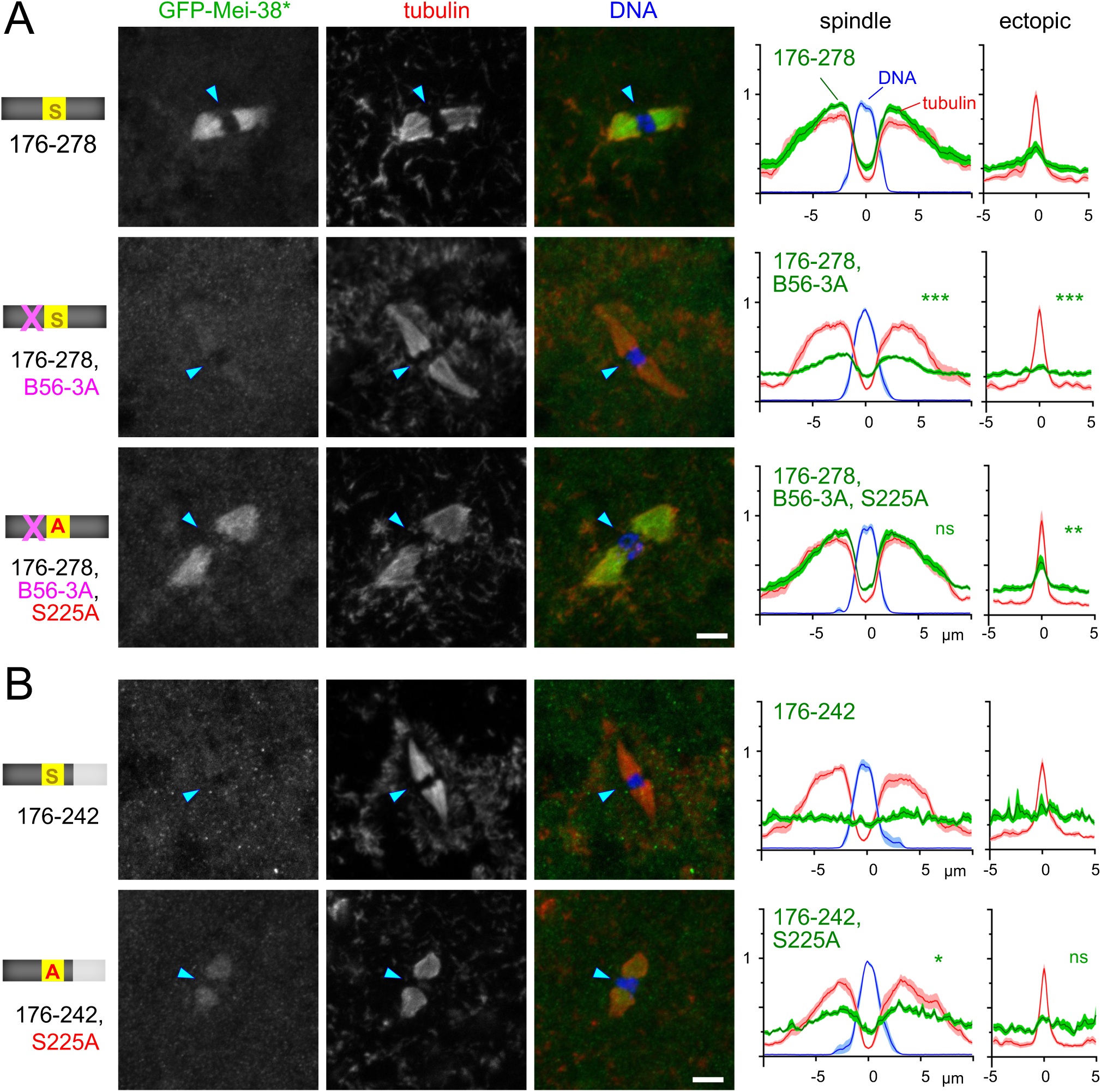
Dephosphorylation of S225 by PP2A-B56 may promote spindle microtubule binding of Mei-38. (A) Localisation of GFP-Mei-38(176-278) carrying mutations in PP2A-B56 docking motif (B56-3A) and/or a non-phosphorylatable mutation at S225 (S225A) in taxol-treated oocytes. The arrowheads indicate the position of the chromosomes. Bar=5 μm. The signal distribution was quantified as Fig.4D. The panel of GFP-Mei-38(176-278) is identical to Fig.4D, and is shown for comparison. ***, ** and ns indicate p<0.001, p<0.01 and p>0.05, respectively, in two-tailed t-tests when the signal intensity of a GFP-Mei-38 variant is compared to that of 176-278.. (B) Localisation of GFP-Mei-38(176-242) with or without a non-phosphorylatable mutation at S225 (S225A) in taxol-treated oocytes. The arrowheads indicate the position of the chromosomes. Bar=5 μm. The signal distribution was quantified as Fig.4D. * and ns indicate p<0.05 and p>0.05, respectively, in two-tailed t-tests when the signal intensity of a GFP-Mei-38(176-242,S225A) is compared to that of 176-242.

We then looked into the role of the C-terminal flanking sequence 243-278. Mei-38(176-242) that lacks this sequence 243-287 has intrinsic microtubule-binding activity but fails to localise to the spindle in oocytes (**Fig 5B**). Addition of this sequence 243-278 enables the spindle localisation in oocytes. We hypothesised that the sequence 243-278 promotes dephosphorylation of Mei-38 on the spindle. Without this sequence, Mei-38(176-242) remains to be fully phosphorylated in oocytes even near the chromosomes/spindle, preventing its binding to both spindle and ectopic microtubules. To test this hypothesis, we mutated S225 of this fragment to alanine. As predicted, this mutant fragment, Mei-38(176-242,S225A), localised to the spindle microtubules in taxol-treated oocytes, although weakly (**Fig 5B)**. Therefore, this fragment, Mei-38(176-242), failed to localise to the spindle microtubules in taxol-treated oocytes, partially because the sequence 243-278 is important for dephosphorylation at S225 which normally takes place in the spindle area.

The S225A mutation abolished spatial regulation of Mei-38(176-278), allowing it to localise equally to both spindle and ectopic microtubules. In contrast, introduction of S225A to either Mei-38(176-278, B56-3A) or Mei-38(176-242) enhanced the localisation to the spindle microtubules, but the localisation to ectopic microtubules was weaker than the spindle. (**Figs 5A and B**). This indicates that, while S225 is solely responsible for spatial regulation of Mei-38(176-278), there is a cryptic mechanism that spatially regulates Mei-38 in the absence of the PP2A-B56 docking site or the C-terminal region 243-278.

Finally, we tested importance of S225 and PP2A-B56 docking motif for the localisation of the full-length Mei-38. We first expressed the GFP-tagged full-length Mei-38 with non-phosphorylatable or phospho-mimetic mutations at S225, Mei-38(S225A) and Mei-38(S225D), in oocytes (**Fig 6A**). The distribution pattern of the non-phosphorylatable mutant, Mei-38(S225A), was similar to Mei-38 without the mutation in taxol-treated oocytes, although the signal appears to be weaker (**Fig 6B**). On the other hand, a phospho-mimetic mutant, Mei-38(S225D), failed to localise to either spindle or ectopic microtubules (**Fig 6B**). This suggests that single phosphorylation of S225 is sufficient to prevent the full-length Mei-38 from binding to microtubules, but the full-length Mei-38 also has another mechanism that suppresses microtubule binding away from the chromosomes, which is consistent to our truncation analysis. Next, to test whether PP2A-B56 docking is important for the localisation of the full-length Mei-38, we expressed the GFP-tagged full-length Mei-38 with mutated PP2A-B56 docking site, Mei-38(B56-3A), in oocytes. The distribution of the Mei-38(B56-3A) localisation in taxol-treated oocytes was similar to Mei-38 without the mutation, but the signal was generally weaker (**Fig 6C)**. This docking site may be less important in the context of the full -length Mei-38.

**Figure 6.**
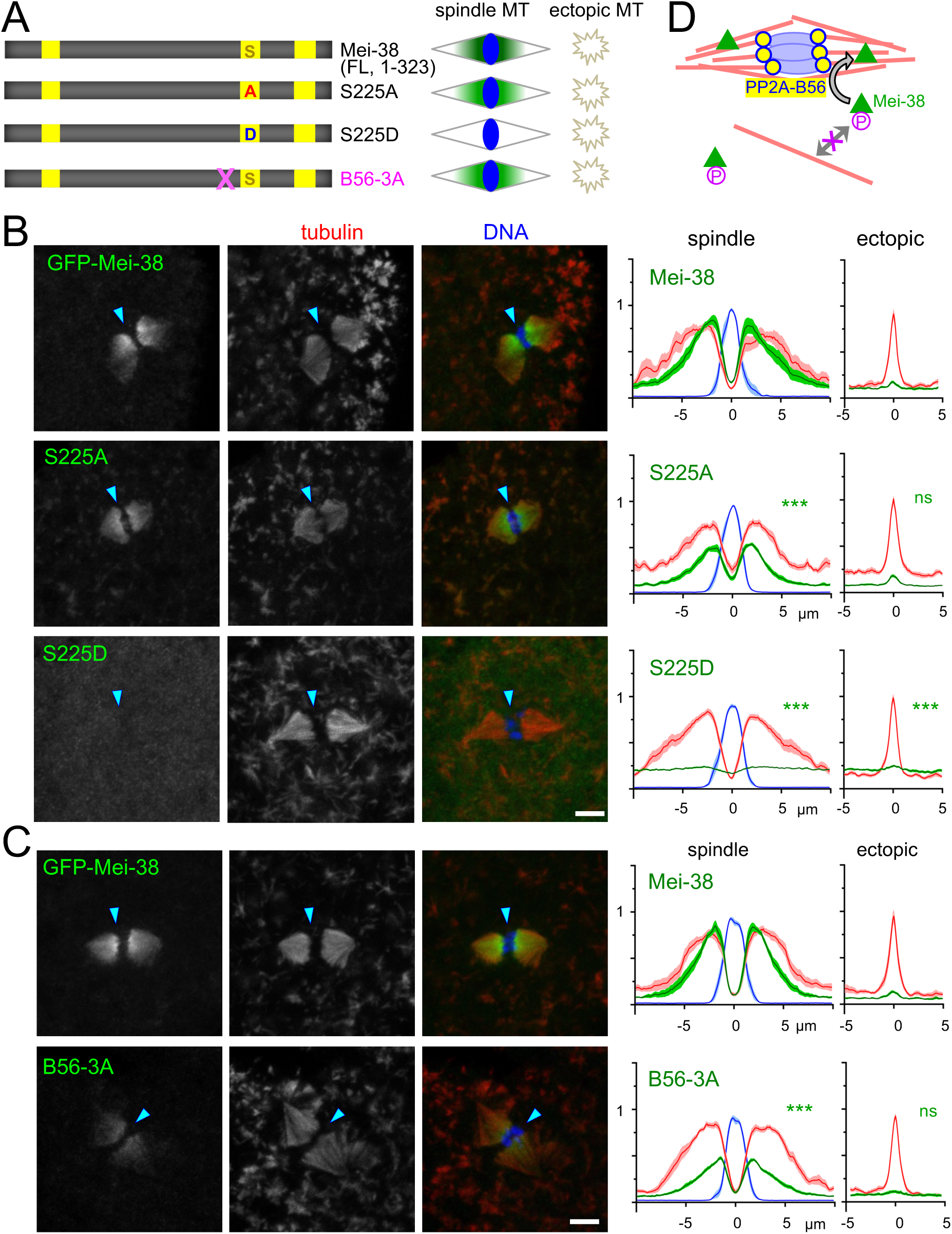
A phospho-mimetic mutation at S225 prevents Mei-38 from binding to microtubules in taxol-treated oocytes. (A) A summary diagram of the GFP-tagged full-length Mei-38 with mutations in S225 and a PP2A-B56-docking motif, together with their localisation in taxol-treated oocytes. Mei-38(S225A) and GFP-Mei-38(S225D) carry non-phosphorylatable and phospho-mimetic residue (A and D) at S225, respectively. (B) Localisation of GFP-Mei-38, GFP-Mei-38(S225A) and GFP-Mei-38(S225D) and GFP-in taxol-treated oocytes. The methods and quantification are the same as Fig.2B. These oocytes were processed and observed in parallel. The arrowheads indicate the position of the chromosomes. Bar=5 μm. (C) Localisation of GFP-Mei-38, and GFP-Mei38(B56-3A) in taxol-treated oocytes. The methods and quantification are the same as Fig.2B. These oocytes were processed and observed in parallel. The arrowheads indicate the position of the chromosomes. Bar=5 μm. (D) A proposed model in which the phosphatase PP2A-B56, concentrated at kinetochores, removes inhibitory phosphorylations from Mei-38, enabling it to bind to spindle microtubules.

## Discussion

The bipolar spindle forms only around the meiotic chromosomes in oocytes without centrosomes in the exceptionally large cytoplasm. This requires key proteins to be locally activated in oocytes. In this study, we have developed a novel method to assess spatial regulation of specific proteins in oocytes by inducing ectopic microtubules using taxol. We have identified 10 spindle-associated proteins that are spatially regulated in *Drosophila* oocytes in terms of microtubule binding. Our analysis of the *Drosophila* TPX2 homologue Mei-38 suggests a phosphatase-driven mechanism, in which inhibitory phosphorylation of Mei-38 is removed by PP2A-B56, concentrated at kinetochores, enabling it to bind to spindle microtubules (**Fig 6D**).

In oocytes, spatially restricted activation of key proteins around the chromosomes is important for the following reasons. (1) Oocytes lack centrosomes, the main microtubule organising centres in mitotic cells, and need to restrict a high level of microtubule nucleation/stabilisation to the proximity of the chromosomes. (2) Non-spindle microtubules co-exist with spindle microtubules in oocytes, but should not organise into bipolar spindles. Key microtubule organising proteins, such as cross-linkers and motors, must be active only near the chromosomes. (3) The density of non-spindle microtubules in oocytes is low, but the volume of the oocyte is far larger than that of the spindle (>1,000 times in *Drosophila* oocytes). A large number of non-spindle microtubules may outcompete spindle microtubules for binding of key microtubule-associated proteins. To overcome this, oocytes need to either produce a far larger amount of the proteins than required, or activate the microtubule-binding activity of these proteins only near the chromosomes.

Understanding the mechanisms governing spatial regulation of proteins important for spindle formation in oocytes remains still limited. For local activation of the spindle-associated proteins in oocytes, Rcc1 (GEF) and the CPC (a kinase complex) are the two chromosomal signals that have been identified so far. Further studies have identified the downstream pathways leading to local activation of key proteins, which involve Ran-Importin and 14-3-3, respectively [4, 5, 28, 32] However, it is still unknown whether they account for all of the spatial regulation in oocytes, as no systematic studies have been carried out to identify chromosomal signals, spatially-regulated proteins or the underlying regulatory mechanisms.

Although proteins regulated by known mechanisms have been systematically identified [33, 34] these studies could not uncover an entirely new mechanism of spatial regulation. Instead, comprehensive understanding of spatially restricted spindle assembly in oocytes requires the systematic identification of spatially regulated proteins regardless of the mechanisms. In-depth studies of each spatially-regulated protein could discover a new chromosomal signal and/or a signaling pathway, or add to a list of proteins controlled by a known mechanism. Studies of all spatially regulated proteins could lead to comprehensive understanding of chromosomal signals and the regulatory mechanisms.

We developed a novel method to visualise a spatial distribution of microtubule binding affinity by inducing ectopic microtubule clusters in *Drosophila* oocytes. Using the method, we have successfully identified 10 spindle-associated proteins that are spatially regulated in terms of microtubule binding. A limitation of our method is that it only assesses general microtubule-binding activity. It does not assess interaction with dynamic microtubules, or other activities, such as nucleation, cross-linking or motor activities. Furthermore, spatial regulation of some proteins may be indirect, as they may bind to another proteins that is spatially regulated or recognise spatially restricted signatures or structures. Nevertheless, our newly developed method is straightforward and rapid, and can be easily applied to other species to uncover conserved principles for spatial regulation in oocytes.

Among spatially regulated proteins we identified, we focused our studies on the *Drosophila* TPX2 homologue Mei-38, as it appear to lack sites potentially regulated by known mechanisms. Our truncation and mutational analysis has identified a microtubule-binding domain (176-278) in Mei-38 that contains a highly conserved sequence among the TPX2 family [30]. We demonstrated the microtubule-binding activity of this domain *in vitro*, in interphase cultured cells and in metaphase I oocytes. Previous studies have identified multiple microtubule-binding domains in vertebrate TPX2 [35–38] including a recent comprehensive study which tested all 10 modules in TPX2 individually and in combinations both *in vitro* and in cells. They detected microtubule binding activity in six modules, but not in the module R6 which possibly corresponds to the microtubule-binding domain we identified in Mei-38. However, it is uncertain which module in TPX2 corresponds to this domain of Mei-38, as the overall organisation and primary sequence differ significantly between the two. All modules in TPX2 may have a potential microtubule-binding activity with considerable redundancy, but the actual activity of each module may change during evolution.

Our analysis of Mei-38 revealed a mechanism possibly driven by dephosphorylation. The spindle-binding domain, Mei-38(176-278), is under spatial regulation in oocytes, showing much stronger localisation to the spindle microtubules than to ectopic microtubules. A phospho-mimetic mutation at S225 prevented binding to any microtubules, while a non-phosphorylatable mutation abolished the spatial regulation, resulting in indiscriminate binding to both spindle and ectopic microtubules. These results suggest that phosphorylation at S225 inhibits microtubule binding, and that this inhibitory phosphorylation is removed exclusively in the spindle area to allow binding to spindle microtubules.

Our results further suggest that the protein phosphatase PP2A-B56 promotes spindle microtubule binding of Mei-38(178-278) by targeting S225 through the adjacent PP2A-B56 docking motif. Additionally, our results also suggest the involvement of another flanking region in dephosphorylation at S225, and the presence of a cryptic mechanism which becomes evident only in the absence of the PP2A-B56 docking site or the flanking region. Furthermore, another region outside of 178-278 also contributes to spatial regulation, suggesting multiple layers of regulation on Mei-38.

As one of the two isoforms of PP2A-B56 is concentrated at the kinetochores in *Drosophila* oocytes [39] we propose that this phosphatase serves a new chromosomal signal that locally activates proteins important for spindle formation (**Fig 6D**). In oocytes, studies on PP2A-B56 have primarily focused on its regulation of kinetochore-microtubule attachment [40–43] A protein phosphatase represents a new type of chromosomal signals, distinct from the previously known signals, a kinase (the CPC) and a GEF (Rcc1). Since inhibitory phosphorylation is a common way to block protein-protein interactions including microtubule binding [44–49] phosphatase-driven mechanisms may be widely used to locally activate more spindle-associated proteins near the chromosomes in oocytes. Furthermore, considering that PP2A-B56 is concentrated at the kinetochores also in mammalian oocytes [50] this phosphatase may act as a chromosomal signal in mammalian oocytes as well. It would be of great interest to test potential roles of PP2A-B56 or other phosphatases in mammalian oocytes.

## Materials and Methods

### Molecular techniques

Standard molecular techniques were followed [51] Plasmids expressing Mei-38 or its variants were generated as below. A Gateway donor vector pENTR was linearized with NotI and AscI. Gene regions of interest flanked by a stop codon and overlapping regions to the ends of the linearised pENTR were amplified from a cDNA from the Nick Brown embryonic library [52]using PrimeSTAR DNA polymerase (Takara) or custom synthesised (IDT). This cDNA (pAG41) encodes a protein (S4 Fig) which is most similar to Mei-38-PA but shorter by two residues. After purification, the linear vector and PCR products were ligated by Gibson assembly (HiFi; New England Biolabs). Sanger sequencing (Genewiz) confirmed that no unwanted mutations were introduced during PCR. Gene regions of interest were then recombined into a Gateway destination vector using LR Clonase II enzyme (ThermoFisher).

To introduce mutations of S225 (S225A, S225D) or PP2A-B56 docking motif (B56-3A; from FKDVTE to AKDATA), upstream and downstream regions of a mutation were separately amplified using primers containing the mutation.

Two destination vectors were used for Gateway cloning. To express genes in flies under the UASp, φPGW was modified from the destination vector pPGW of the Murphy’s Gateway collection by inserting the φC31 attB recombination sequence at the AatII site. To express genes under the Cu^2+^-inducible metallothionein promoter in *Drosophila* S2 cells, the destination vector pMTGW (a gift from G. Goshima, Nagoya University, Japan) was used. To make transgenic flies, expression clones were microinjected into the fly line (BDSC_9750; PBac{y^+^-attP-3B}VK00033) using φC31 integrase-mediated transgenesis, which was carried out by BestGene Inc.

### *Drosophila melanogaster* genetic techniques

Standard *Drosophila* techniques [53] were followed. To test the localisation of an endogenous protein using an antibody, *w^1118^* was used as the wild type. To the localisation of an exogenous GFP-tagged protein in oocytes, virgin females of the *GAL4*-driver line *P[GAL4::VP16-nos.UTR]MVD1* (BDSC_4937) was crossed with males carrying a transgene of the GFP-tagged protein under the control of *UASp*. Transgenic lines with various GFP-tagged spindle-associated proteins were previously reported [54].

### Taxol treatment and immunostaining of *Drosophila* oocytes

Newly eclosed female adult progeny (<1 day old) were matured at 25°C for three days with males and standard cornmeal medium supplemented with dried yeast powder. Ovaries from 24 mature females were dissected in buffer containing 80 mM HEPES pH 6.8, 1 mM MgCl_2_, 1 mM EGTA and 20 μM taxol (T7402; Merck). After incubation in the same taxol-containing buffer for a further 15 minutes, the excess buffer was removed, and then 30 ml of fresh methanol were added.

These methanol-fixed ovaries were used for immunostaining. The ovaries were sonicated with a microprobe using VibraCell (VCX500; Sonics) for a 1-second burst and mature oocytes without the chorion and the vitelline membrane were collected and kept in fresh methanol. Sonication may be repeated to collect enough oocytes. To rehydrate, collected oocytes were incubated in a 400 μl of PBS/methanol (1:1) for 10 minutes, followed by incubation in 400 μl of 100% PBS for 10 minutes. The oocytes were incubated with 100 μl of the blocking buffer (PBS, 0.1% Triton X-100, 10% FBS) for 30 minutes. The oocytes were incubated for 4 hours at 22°C or overnight at 4°C with primary antibodies diluted in 100 μl of the blocking buffer. After washing three times in 200 μl PBST(PBS containing 0.1% Triton X-100) for 10 minutes each, the oocytes were incubated for 2 hours at 22°C with 100 μl of secondary antibodies plus 0.5 mg/ml DAPI in PBST. This was followed by four 15-minute washes in PBST and one in PBS. They were mounted in the mounting medium (85% glycerol and 2.5% propylgallate) between a coverslip and a glass slide sealed with nail varnish.

Antibodies were used for immunostaining as below: anti-α-tubulin (mouse monoclonal DM1A; 1:250; Sigma-Aldrich), anti-TACC (rabbit polyclonal against D-TACC-CTD; 1:1,000; [55]), anti-Ncd (rabbit polyclonal against the full-length, 1:1,000), anti-GFP (rabbit polyclonal; A11122; 1:125; Thermo Fisher Scientific), anti-Mink (rat polyclonal; 1:500; [56], anti-Aurora B (rat polyclonal; 1:1000). Secondary antibodies conjugated with Alexa Fluor 488, Cy3, and Cy5 were used (1:100-1:200; The Jackson Laboratory or Molecular Probes).

After immunostaining, *Drosophila* oocytes were imaged using an Axiovert Z1 microscope (Zeiss) attached to a confocal laser scanning head LSM800 (Zeiss) before 1st October 2024 or an AxioObserver 7 (Zeiss) attached to a LSM900 (Zeiss) after this date. Oocytes were visualised under a Plan-Apochromat objective lens (63×/1.4 numerical aperture) with Immersol 518F oil (Zeiss). Images were captured at 512x512 pixels (100 nm x 100 nm pixel size) and at 0.5 μm Z-intervals. AxioObserver 7 with LSM900 produced similar, but not quantitatively identical, results compared to Axiovert Z1 with LSM800, including lower background signals. Only the images acquired by the same microscope were compared, and presented in the same figure. Data presented in Figs 1, 2, S1-S3, S5 were acquired by AxiovertZ1 with LSM800, while data presented in Figs 4-6 were acquired by AxioObserver7 with LSM900. The images are presented after maximum-intensity projection and a linear brightness enhancement.

To quantify the distribution of GFP-Mei-38, the signal intensities of α-tubulin, GFP and DNA were measured in taxol-treated oocytes (**S8 Table**). After a maximum-intensity projection, a line with 1 μm width was drawn along the long axis of each spindle and an ectopic microtubule cluster with a similar maximum tubulin intensity in the same oocytes, and the intensity (the mean of each 1-μm width) was measured in 0.0978 μm intervals along the lines using ImageJ. For a line for ectopic microtubules, the position of the highest tubulin intensity was set to 0. For a spindle line, the position of the centre of the chromosomal mass was set to 0. When this position was different from the lowest tubulin signal, the mid-point between the two was set to 0. Then, each intensity value was normalised by dividing the maximal value on the line of the spindle in the same oocyte. The graphs show the mean intensity with the standard error along the long axis of the spindle and the line across an ectopic microtubule cluster. When the profiles are compared, each profile of the Mei-38 mean signal intensity is presented after normalisation to make the mean background signal intensity (at 0 in the spindle line) identical to the control. For a two-tailed t-test, the Mei-38 signal intensity on the spindle or ectopic microtubules was compared to the control (at the top row of a figure) at the position where the mean intensity is the highest along the spindle or ectopic line in the control.

### Immunoblotting of *Drosophila* ovaries

Twenty four pairs of ovaries were dissected from mature females in absolute methanol. Methanol-immersed ovaries were washed with PBS once and then 200 μl of PBS was added. After ovaries were homogenised with a pestle, 200 μl of 2x SDS loading buffer (50 mM Tris-Cl pH 6.8, 2% SDS, 10% glycerol, 0.1% bromophenol blue) plus 5% β-mercaptoethanol were added. The mixture was boiled at 95°C for 3 minutes, and 10 μl was loaded on each lane of SDS PAGE gel (Mini Protean Tetra Biorad) for electrophoresis at 150 voltages for 1 hour. Then, proteins were transferred from the gel onto nitrocellulose membranes (Protran 0.2 NC; GE Healthcare) under 100 voltages for 20 minutes. The membrane was stained with a reversible Protein Stain kit (Thermo Fisher Scientific). After destaining, it was incubated with PBSTw (PBS+0.1% Tween 20) containing 3% skim milk for 1 hour. Then it was incubated with the primary antibodies in PBSTw (PBS+0.1% Tween 20) containing 3% skim milk at 22°C overnight with rotation. After three 10-minute washes in PBSTw (0.1% Tween 20), the membrane was incubated with fluorescent secondary antibodies diluted in PBSTw at 22°C for 2 hours with rotation. After four 15-minute washes in PBSTw, the membrane was visualised on an Odyssey CLx imaging scanner (v3.0.30; LI-COR) with 600ppi.

The primary antibodies for immunoblotting were anti-GFP (rabbit polyclonal; A11122; 1:1000; Thermo Fisher Scientific) and anti-α-tubulin (mouse monoclonal DM1A; Sigma-Aldrich; 1:2000). The secondary antibodies used were IRDye 800CW-conjugated goat anti-rabbit (1:20,000) (LI-COR Biosciences, 926-32211), IRDye 800CW-conjugated goat antirat (1:20,000) (LI-COR Biosciences, 926-32219), IRDye 680LT-conjugated goat anti-mouse (1:15,000) (LI-COR Biosciences, 926-68020).

### Live imaging of *Drosophila* Schneider S2 cells

S2 cells were cultured in Schneider media (Invitrogen, 21720024) supplemented with 10% heat-inactivated fetal calf serum (Invitrogen, 10500064) and 1% antibiotics (Sigma, A5955) at 27°C. To observe the localisation of GFP-Mei-38 and its variants, expression plasmids containing inserted genes were transfected using X-tremeGENE HP (Merck). After 24 hours, images of live transfected cells were obtained using an Axiovert microscope (Zeiss) attached to a spinning-disc confocal head (CSU-X1; Yokogawa) controlled by Volocity (PerkinElmer).

### Protein production and purification

To generate the antigen of Mei-38 antibody, we produced N-terminal maltose binding protein tagged Mei-38 (1-278). The entry plasmid pENTR-Mei-38(1-278) was recombined using LR Clonase with a destination vector containing the Gateway cassette at the XmnI site of pMALc2. *E. coli* (BL21(DE3)/pLysS) carrying the plasmid was cultured to saturation at 37°C and diluted 1:100 to 1000 ml of LB with ampicillin and chloramphenicol. After 2 hours at 37°C, a final 1 mM IPTG was added to the culture. After further culturing at 37°C for 3 hours, bacteria were centrifuged and re-suspended in 5 ml Lysis buffer (50 mM Tris-HCl pH8, 1 mM EDTA, 100 mM NaCl) and lysed by repeated freeze-thaw cycles on dry ice and water baths until the cell suspension became viscous. After adding a final 250 μg/ml lysozyme, the lysate was incubated on ice for 10 minutes, then at 37°C for 10 minutes, followed by supplementing with final 10 μg/ml DNase and 2 mM MgCl_2_. It was left at 37°C for 10 minutes or longer until the viscosity decreased. Five ml PBS was added to the lysate, which was then cleared by centrifugation at 14,000 rpm using Eppendorf FA-45-18-11 for 10 minutes at 4°C. The supernatant was applied to a column containing 1 ml of amylose resin (NEB) prewashed by 5 ml of PBS, and the flowthrough was collected. After washing twice with 20 ml of PBS, the MBP-tagged proteins were then eluted through a column by adding a PBS containing 10 mM maltose. Five fractions of 500 μl were collected, and then 50 μl of each fraction was mixed with equal volume of 2x SDS loading buffer and final 5% β-mercaptoethanol. Samples were boiled for 3 minutes at 95°C and analysed by SDS-PAGE with BSA for quantification. The first three eluted fractions from two rounds of purification were mixed together and then concentrated using the centrifugal filter (Merck Millipore) to get a final volume of 2 ml of 0.2 mg/ml. Two rats were immunised four times each by Eurogentec using 50 μg of the purified protein each.

To perform microtubule-binding assay *in vitro*, we produced Mei-38(176-242) with a maltose binding protein (MBP) tag at the N-terminus. The entry plasmid pENTR-Mei-38(176-242) was recombined using LR Clonase with the same destination vector as MBP-Mei-38(1-278). MBP-Mei-38(176-242) was expressed and purified following the same procedure as MBP-Mei-38(1-278), except 500 ml of bacteria culture and 3 ml lysis buffer were used (**S7 Fig)**.

### *In vitro* microtubule-binding assay

To test the microtubule-binding activity of MBP-Mei-38 (176-242aa), 10 μl of 5 mg/ml pure tubulin from porcine brains (T240: Cytoskeleton Inc) in BRB80 (80 mM HEPES pH 6.8, 1 μM MgCl_2_, 1 μM EGTA) containing 1 μM GTP was diluted with 40 μl of BRB80 Buffer. 1.8 μg of freshly made MBP-Mei-38 (176-242) protein was mixed with diluted tubulin before addition of 10 μg BSA protein, and then BRB80 buffer up to a total volume 200 μl. After centrifugation at 22°C with 45,000 rpm using a Beckman TLA120.2 rotor for 20 minutes to remove insoluble proteins, the supernatant was split into two fractions of 50 μl each. To one fraction (sample), final 20 μM paclitaxel (T7402 Merck) and 1 μM GTP were added, and the same volume of BRB80 was added to the other (control). Both samples were incubated at 22°C for 10 minutes and then centrifuged at 22°C at 14,000 rpm using an Eppendorf FA-45-18-11 rotor for 15 minutes to pellet microtubules. The sample pellet was washed with 100 μl of BRB80 containing 20 mM paclitaxel and 1 μM GTP and the control pellet was washed with 100 μl of BRB80. After the pellets were resuspended in 25 μl of BRB80, 25 μl of 2x SDS sample buffer and final 5% β-mercaptoethanol were added to the supernatants and pellets. They were boiled and 10 μl were loaded on each lane for western blot using an anti-MBP antibody (rat polyclonal against the full length; 1:1,000) as above.

### Data and resource availability

All reagents generated in this study, including fly lines and plasmids, are available on request. All relevant data and details of resources can be found within the article and its supplementary information.

## Acknowledgements

We are grateful to the current and past members of the Ohkura laboratory for their help and contributions, and especially to Fiona Cullen for initially developing the taxol assay. Stocks from the Bloomington Drosophila Stock Center (National Institutes of Health grants P40OD018537) were used in this study. This work is supported by research grants to HO from the Wellcome Trust (206315, 227907). A part of the work was carried out in the former Wellcome Centre for Cell Biology supported by core funding from the Wellcome Trust (203149). XW received a PhD studentship from Darwin Trust of Edinburgh. The authors declare no competing financial interests.

## Supporting information

**S1 Fig.**
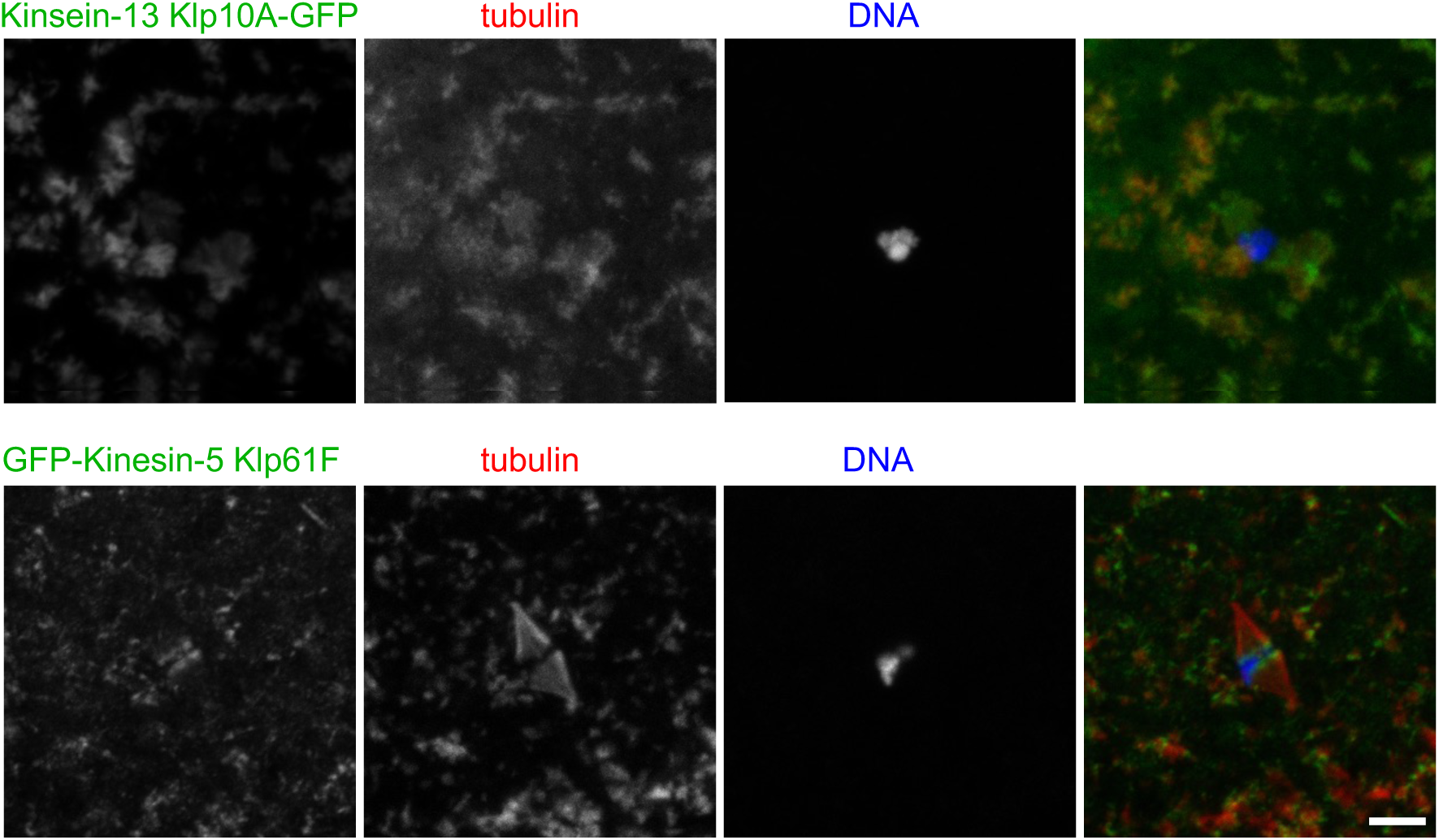
Spindle proteins that are not regulated spatially in oocytes in terms of microtubule binding. Mature oocytes expressing a GFP-tagged protein were incubated with taxol and immunostained using GFP and α-tubulin antibodies. GFP-tagged Kinesin-13 Klp10A and Kinesin-5 Klp61F localised to both spindle and ectopic microtubules with similar intensities. Bar=5 μm.

**S2 Fig.**
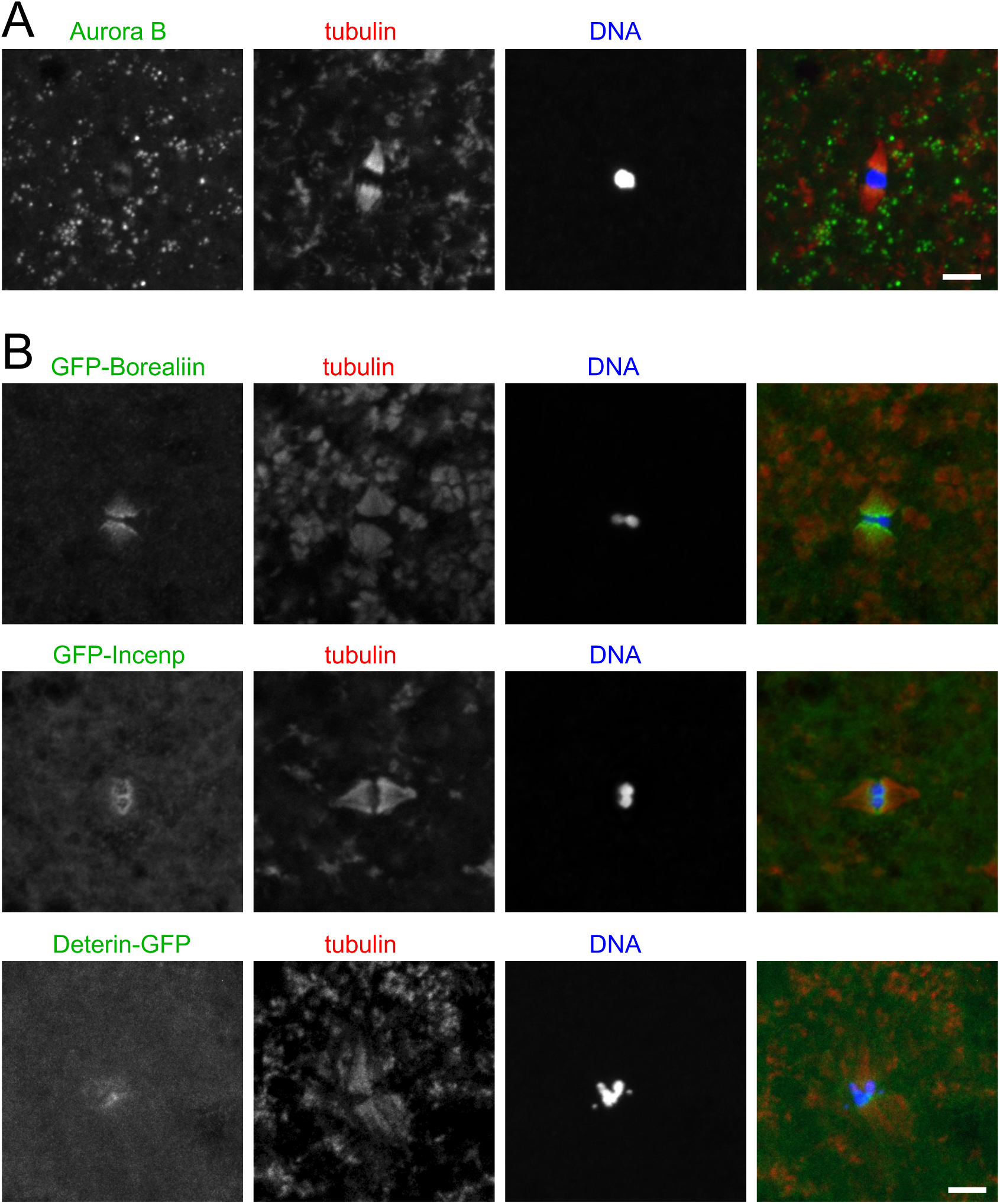
Microtubule binding of the CPC subunits are spatially regulated in oocytes. (A) Wild-type mature oocytes arrested in metaphase I were incubated with taxol and immunostained using Aurora B and α-tubulin antibodies. Aurora B was concentrated on spindle microtubules near chromosomes, not on ectopic microtubule clusters. Bar=5 μm. (B) Mature oocytes expressing a GFP-tagged protein were incubated with taxol and immunostained using GFP and α-tubulin antibodies. GFP-tagged Borealin, Incenp and Deterin were concentrated on spindle microtubules near the chromosomes, not on ectopic microtubule clusters. Bars=5 μm.

**S3 Fig.**
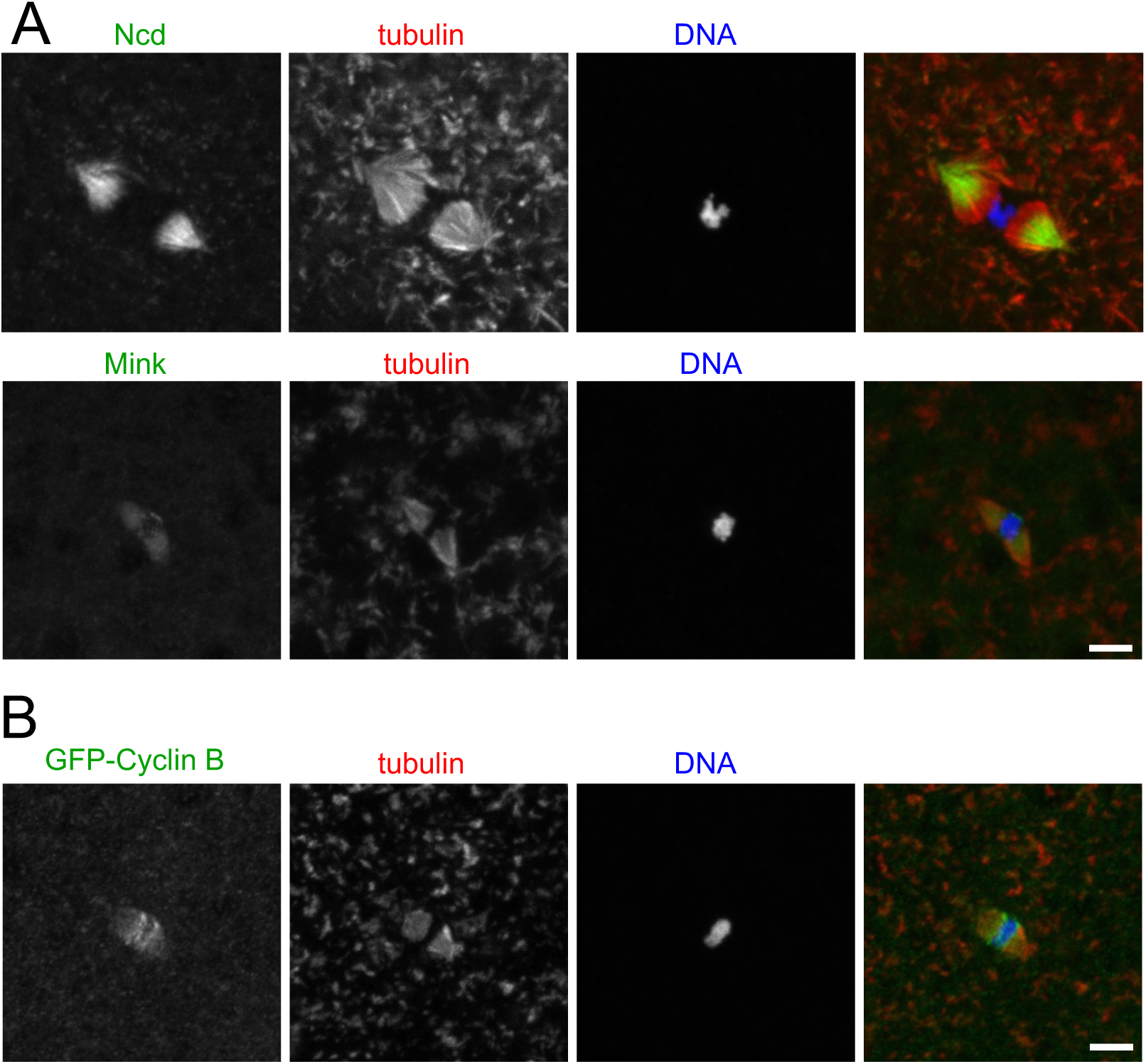
Microtubule binding of Ncd, Mink, Cyclin B and Mars are spatially regulated in oocytes. (A) Wild-type mature oocytes arrested in metaphase I were incubated with taxol and immunostained using antibodies against Ncd or Mink and α-tubulin. Ncd and Mink were concentrated on spindle microtubules, but much less on ectopic microtubule clusters. Bar=5 μm. (B) Mature oocytes expressing a GFP-tagged Cyclin B were incubated with taxol and immunostained using antibodies against GFP and α-tubulin. GFP-tagged Cyclin B was concentrated on spindle microtubules near chromosomes, not on ectopic microtubule clusters. Bar=5 μm.

**S4 Fig.**
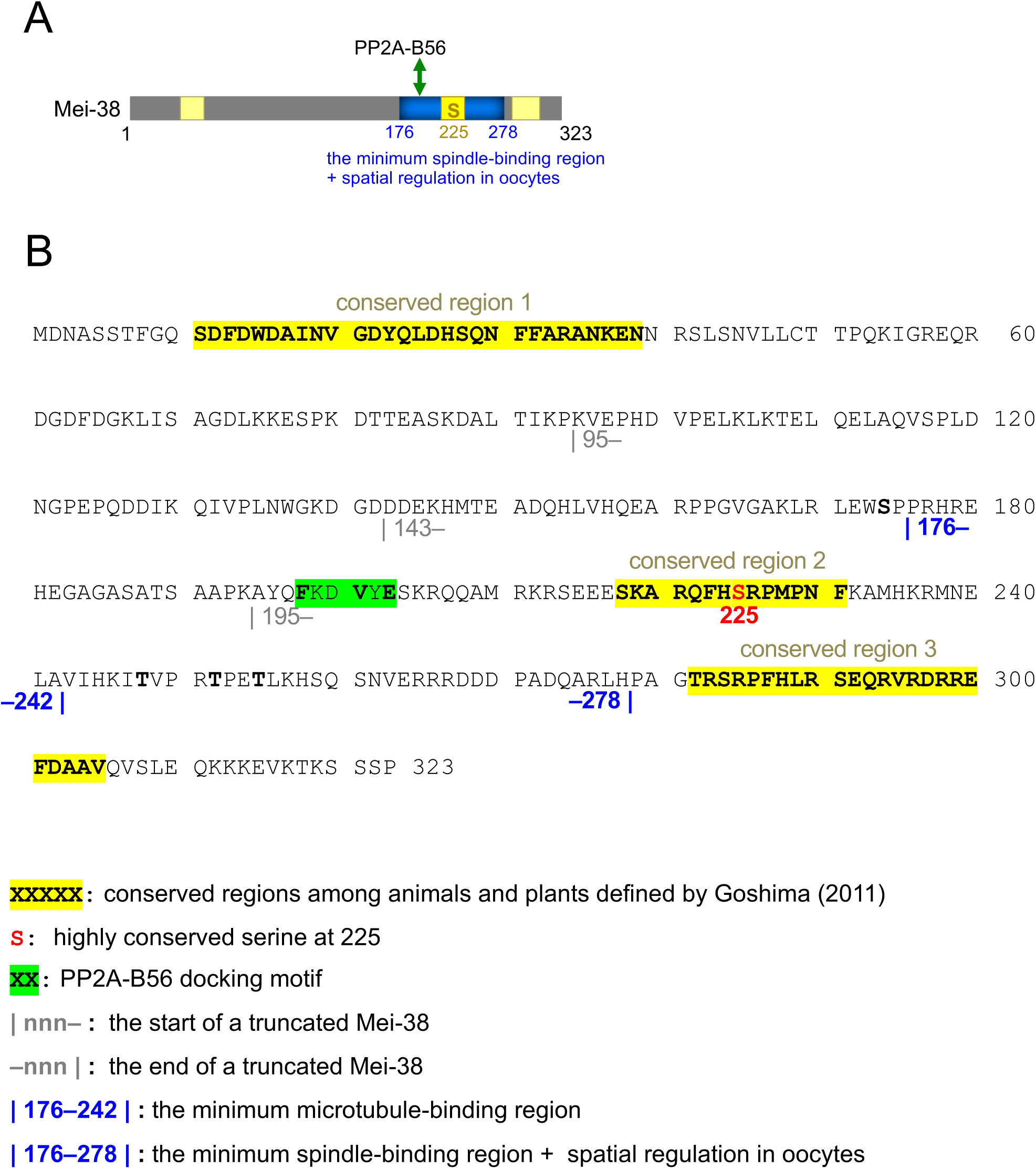
The Mei-38 amino acid sequence. (A) Schematic diagram of the Mei-38 primary structure. (B) The Mei-38 amino acid sequence based on the sequence of the cDNA pAG41. Key residues are marked. Bars=5 μm.

**S5 Fig.**
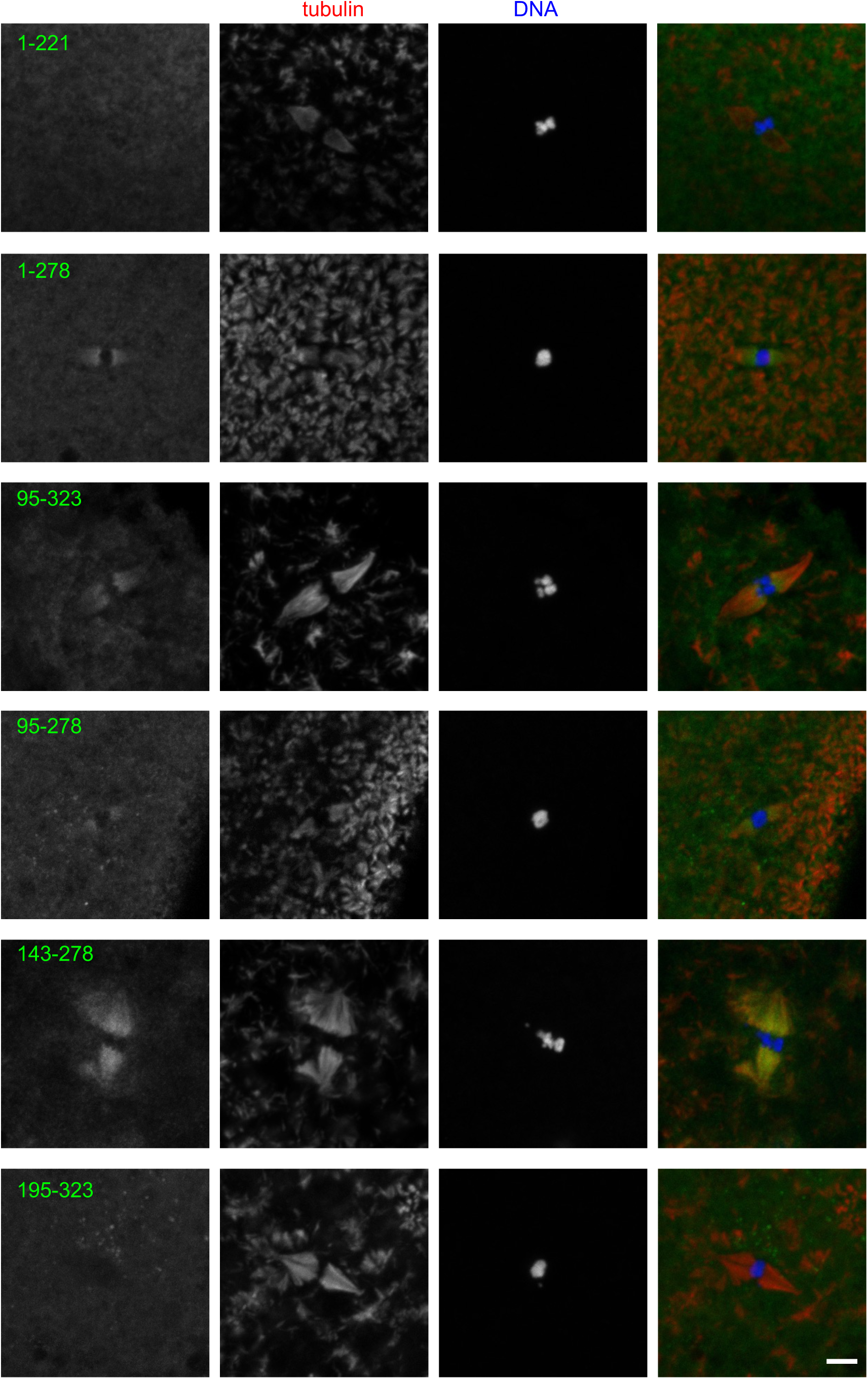
Localisation of various Mei-38 truncations in taxol-treated oocytes. After taxol treatment, mature oocytes expressing various GFP-tagged Mei-38 truncations were immunostained using GFP and α-tubulin antibodies. Quantification of these truncations and the images and quantification of other truncations are shown in Fig.2. Bar=5 μm.

**S6 Fig.**
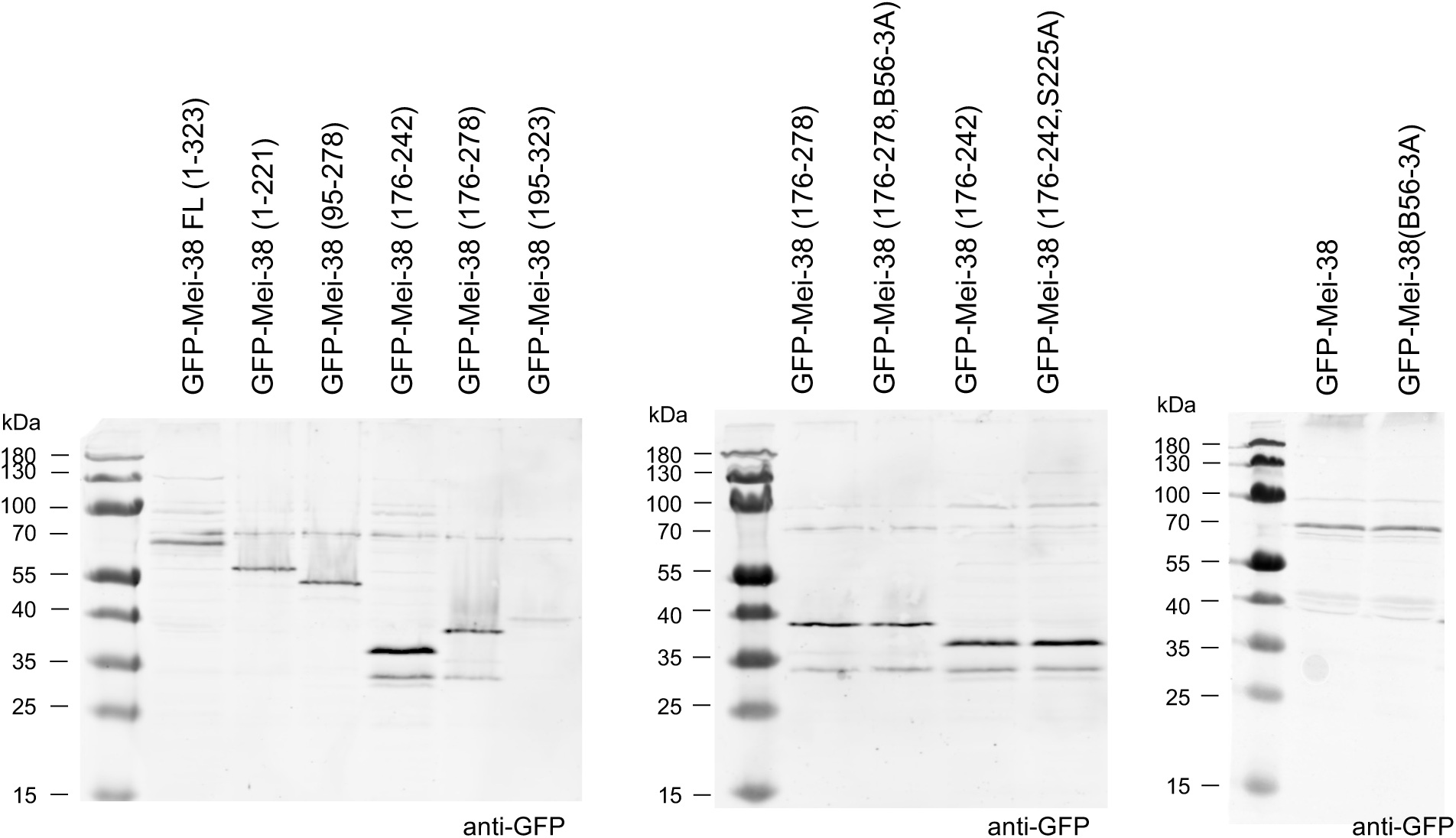
Western blots of various GFP-Mei-38 truncations and mutations. Ovaries expressing various forms of GFP-Mei-38 were run on SDS-PAGE, and western-blotted using an anti-GFP antibody to estimate the size and quantity of GFP-tagged proteins. Comparable amounts of GFP-tagged proteins were produced except Mei-38(195-323).

**S7 Fig.**
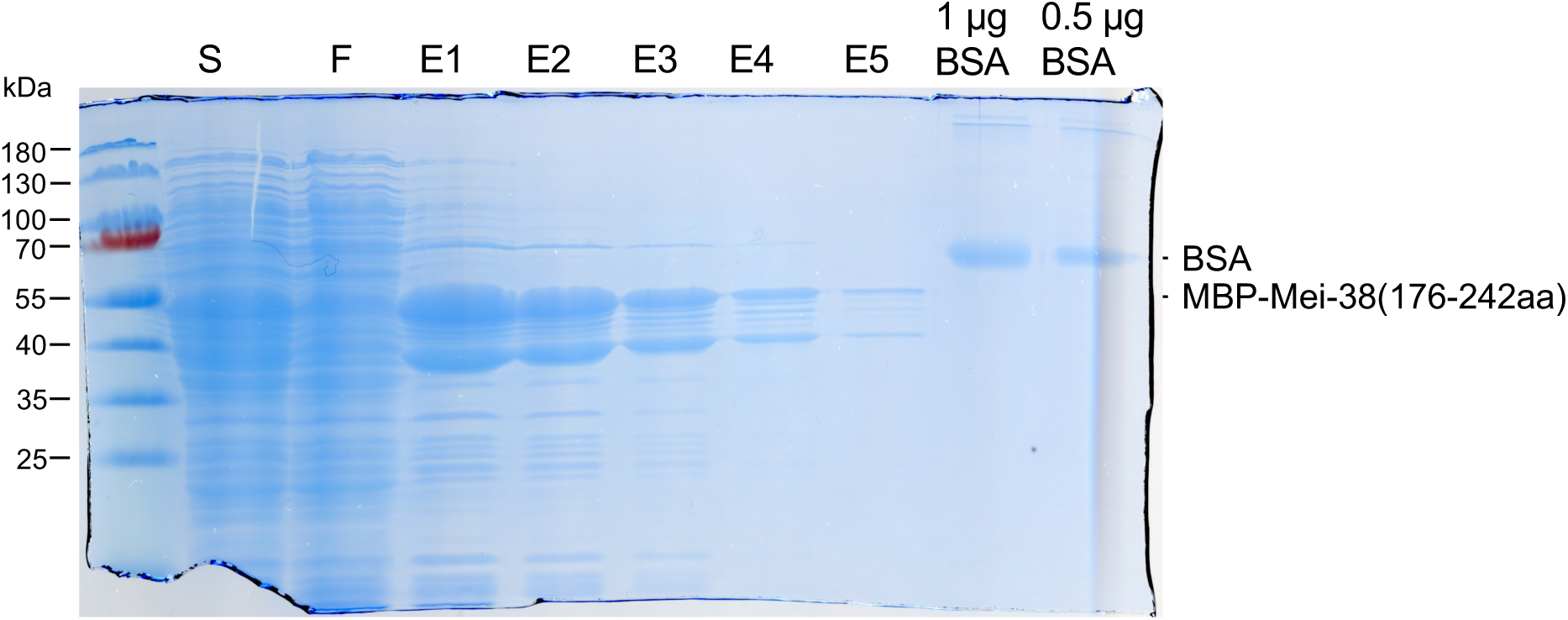
Purification of MBP-Mei-38(176-242) After bacterial cells expressing MBP-Mei-38(176-242) were lysed, the lysate was centrifuged and the supernatant (S) was run through an amylose column. The flowthrough fraction (F) and elution (E1-E5) using buffer containing maltose were analysed together with BSA for quantification.

**S8 Table.** Measurements of the signal intensities. An Excel file containing all measurements of signal intensities used for producing graphs in the figures.

